# Deletion of wheat alpha-gliadins from chromosome 6D improves gluten strength and reduces immunodominant celiac disease epitopes

**DOI:** 10.1101/2024.07.19.604379

**Authors:** Maria G. Rottersman, Wenjun Zhang, Junli Zhang, Gabriela Grigorian, German Burguener, Claudia Carter, Teng Vang, Joshua Hegarty, Xiaoqin Zhang, Jorge Dubcovsky

**Affiliations:** Dept. of Plant Sciences, University of California, One Shields Avenue, Davis, CA, 95616, USA; Proteomics Core Facility, University of California, 451 E. Health Sciences Dr., Davis, CA 95616, USA; California Wheat Commission, 1240 Commerce Ave., Woodland, CA, 95776, USA; Howard Hughes Medical Institute, 4000 Jones Bridge Rd, Chevy Chase, MD, 201815, USA

**Keywords:** wheat, celiac disease, prolamins, gliadins, gluten, breadmaking quality, deletions

## Abstract

Wheat gliadins and glutenins confer valuable end-use characteristics but include amino acid sequences (epitopes) that can elicit celiac disease (CeD) in genetically predisposed individuals. The onset of CeD in these individuals is affected by the amount and duration of the exposure to immunogenic epitopes. Therefore, a reduction of epitopes that result in high immune responses in the majority of CeD patients (immunodominant epitopes) can reduce the incidence of CeD at a population level. We identified deletions encompassing the α-gliadins at the three wheat genomes, designated hereafter as *Δgli-A2* (PI 704906), *Δgli-B2* (PI 704907), and *Δgli-D2* (PI 704908). The *Δgli-D2* deletion, which eliminates major immunodominant epitopes, significantly increases gluten strength, improves breadmaking quality, and has no negative effects on grain yield or grain protein content. By contrast, *Δgli-A2* and *Δgli-B2* showed limited effects on breadmaking quality. The stronger effect of the *Δgli-D2* deletion on gluten strength is associated with the presence of α-gliadins with seven cysteines in *GLI-D2* that are absent in *GLI-A2* and *GLI-B2* loci, which all have α-gliadins with six cysteines. We show that α-gliadins with seven cysteines are incorporated into the gluten polymer, where they likely function as chain-terminators limiting the expansion of the gluten polymer and reducing its strength. In summary, the publicly available *Δgli-D2* deletion developed in this study can be used to simultaneously improve wheat gluten strength and reduce immunodominant CeD epitopes.

## 1. Introduction

Celiac disease (CeD), a common immune-mediated condition characterized by inflammation of the small intestine, affects approximately 1% of the world’s population (Bradauskiene et al. 2023), and its incidence has been increasing over time (King et al. 2020). CeD is triggered in genetically predisposed individuals by long term consumption of wheat gluten and gluten-like peptides. Currently the only fully effective treatment for CeD is a completely gluten-free diet. However, even those who strictly adhere to a gluten-free diet may suffer from compromised health due to nutritional imbalance and a less robust gut microbiome, social isolation, and an increased cost of living (Sánchez et al. 2023).

Gluten is a protein polymer that confers wheat bread and pasta dough its viscoelastic and extensible properties. This complex includes two classes of proteins: glutenins, which are interconnected to each other by disulfide bonds at their cysteine residues (CYS) forming a mesh-like structure, and gliadins, which typically form non-covalent bonds with glutenins (Shewry et al. 2022). Most glutenin and gliadin proteins include amino acid sequences (epitopes) that can bind to specific human leukocyte antigen (HLA) receptors eliciting an immune response in people with CeD. This binding is recognized by gluten-sensitive T-cell lymphocytes resulting in cytokine release and damage to the intestinal lining (Vader et al. 2002). Epitopes that result in consistently high immune responses in the majority of CeD patients are designated as immunodominant epitopes (e.g., DQ2.5-glia-α1, DQ2.5-glia-α2, DQ2.5-glia-ω1, and DQ2.5-glia-ω2) (Jabri et al. 2017; Tye-Din et al. 2010).

Multiple studies with children, including some with identical twins, have shown that the amount of consumed immunogenic wheat epitopes is a factor correlated with the incidence of CeD (Ivarsson et al. 2000; Mearin et al. 1983). In addition, people that carry the HLA-DQ8 have a lower risk of developing CeD than people carrying the HLA-DQ2 allele, which recognizes a larger number of epitopes than HLA-DQ8 (Megiorni et al. 2009). These results suggest that the quantity of ingested immunogenic epitopes may affect the onset of the disease (Koning 2012), and that a reduction of the immunogenic epitopes present in wheat may reduce the incidence of the disease at a population level.

However, elimination of all wheat immunogenic epitopes is a complex task since there are more than 40 unique CeD-toxic proteins currently known in the bread wheat genome (Chlubnova et al. 2023; Sollid et al. 2012; Sollid et al. 2020). Moreover, elimination of all the glutenin and gliadin proteins carrying CeD epitopes would result in unacceptable reductions in breadmaking quality. The elimination of immunogenic proteins with a limited contribution to breadmaking quality represents a practical intermediate step in the reduction of wheat CeD toxicity.

The α-gliadins, located on the short arms of the chromosomes of homeologous group 6, are an attractive target for this initial step for a number of reasons. First, α-gliadins are mostly monomeric proteins that typically do not form covalent bonds with the gluten polymer and are expected to have a limited effect on gluten strength. However, there are conflictive results on the effect of the deletion of α-gliadins on breadmaking quality (Li et al. 2018; van den Broeck et al. 2009). In addition, α-gliadins account for the greatest number of gliadins in the genomes of pasta wheat Svevo (68%) (Maccaferri et al. 2019) and bread wheat Fielder (55%) (Sato et al. 2021), and are the most abundantly expressed gliadins in the developing grains (Wang et al. 2017). More importantly, the α-gliadins are particularly rich in known immunogenic nanomers (Chlubnova et al. 2023; Sollid et al. 2012; Sollid et al. 2020), including larger peptides with multiple immunodominant epitopes (Anderson et al. 2000; Jabri and Sollid 2017; Shan et al. 2002).

We used gamma radiation to generate deletions encompassing the α-gliadins at the three wheat genomes, which are designated hereafter as *Δgli-A2* (PI 704906), *Δgli-B2* (PI 704907), and *Δgli-D2* (PI 704908). These lines have been deposited in GRIN-global and are available without restrictions. In this study, we characterize the size and genes included in these deletions and their effect on grain yield, grain protein content and breadmaking quality. Finally, we present proteomic results that explain the beneficial effects of the *Δgli-D2* deletion on breadmaking quality relative to *Δgli-A2* and *Δgli-B2* deletions.

## 2. Materials and Methods

### 2.1. Plant materials used in this study

The mutagenized line (RIL143, *Triticum aestivum* L. subsp. *aestivum*) is an F_7_-derived F_8_ recombinant inbred line developed via single-seed descent from the cross between UC1110 and PI 610750. UC1110 is a hard white spring breeding line from the University of California, Davis (GRIN-Global accession GSTR 13501, pedigree Chukar/3/Yding//Bluebird/Chanate), and PI 610750 is a synthetic derivative from CIMMYT (PI 610750, pedigree: Croc1/*Aegilops tauschii* (Synthetic 205)//Kauz) (Lowe et al. 2011). RIL143 carries the adult plant stripe rust resistance gene *Yr34* (synonym *Yr48*) from PI 610750 and was mutagenized with gamma radiation as part of the efforts to fine map this resistance gene (Chen et al. 2021). We originally treated 2,076 grains with 300 GY of radiation from which we obtained 850 fertile M_1_ plants. M_2_ head rows were grown in the field at UC Davis in 2012-2013, and 650 M_3_ head-rows were grown in the field in 2013-2014, from which we harvested single spikes of M_4_ seeds which were used in this study.

### 2.2. Haplotype analysis

To select the best genome reference for each of the RIL143 α-gliadin loci, we performed a haplotype analysis. We used SNPs from the α-gliadin genes and ten flanking genes on both sides of the gliadin region, extracted from the publicly available WheatCAP exome capture data in T3 (Blake et al. 2016) (https://wheat.triticeaetoolbox.org/downloads/download-vcf.pl, genotype trial “Exome_Capture_2017_WheatCAP_UCD”). Based on Chinese Spring Refseq v1.0 coordinates, the SNPs were extracted from chromosome 6A between 24,073,547 and 26,530,524 bp, from chromosome 6B between 42,773,234 and 45,164,040 bp and from the unassigned chromosome between 93,797,403 and 94,934,120. These genes are in the same scaffold, which corresponds to the D genome in Fielder and other sequenced genomes. Lines with more than 50% of missing data were excluded from analysis. We performed a cluster analysis using R functions ‘dist’ and ‘hclust’ (method = ‘ward.D2’)(R-Core-Team 2021). Haplotypes were determined using 30% of the maximum clustering height.

After the haplotype analysis, we extracted 50 bp sequences flanking the informative SNPs and blasted them against 20 sequenced wheat genomes (Athiyannan et al. 2022; Kale et al. 2022; Maccaferri et al. 2019; Sato et al. 2021; Walkowiak et al. 2020; Zhu et al. 2019) and 5 *Aegilops tauschii* genomes (Luo et al. 2017; Zhou et al. 2021). SNPs were called based on the BLASTN results, combined with the previous SNPs and used to repeat the clustering analysis using the same method as described above.

### 2.3. Identification and characterization of the deletion lines

To identify deletions in α-gliadin gene clusters on the short arms of chromosome 6A, 6B, and 6D, we screened the RIL143 radiation mutants with genome specific primers described in Data S1 and selected the ones with no PCR amplification. Final selected genotypes were verified using SDS-PAGE (Singh et al. 1991). We also developed primers to track a secondary deletion identified in chromosome 6A (primers Gap2F1 and Gap2R1, Data S1), and to confirm that this deletion was separated from *Δgli-A2* in BC_4_F_2_ plants selected from the introgression of *Δgli-A2* into the variety Fielder. Finally, we intercrossed the RIL143 *Δgli-A2* and *Δgli-D2*, and in the progeny selected a line homozygous for both deletions, designated hereafter *Δgli-A2 Δgli-D2*. Double deletions including *Δgli-B2* were sterile.

To test the effect of the *Δgli-D2* deletion in a genetic background with higher yield potential and better breadmaking quality than RIL143, we crossed the *Δgli-D2* deletion in RIL143 with the commercial variety UC-Central Red (PVP 2019-00011). We then backcrossed the F_1_ four more times to the adapted variety and in the progeny selected BC_4_F_2_ plants homozygous for *Δgli-D2.* The linked codominant simple sequence repeat marker BARC54 (Data S1) was used to trace the deletion during the marker assisted selection process. UC-Central Red is a high-yielding hard red spring wheat variety released by the University of California in 2021 that has excellent breadmaking quality, as documented by its inclusion in the list of recommended varieties based on breadmaking quality generated by the California Wheat Commission.

To characterize the size of the deletions, we generated exome capture data using the assay described in (Krasileva et al. 2017). Reads from the wildtype and mutant lines were mapped at high stringency (no SNPs allowed) to the Fielder reference genomes (v1.0) (Sato et al. 2021) using the BWA-MEM algorithm (Li 2013). High stringency mapping was used to minimize errors in the delimitation of the deletion borders generated by reads from close homeologs outside the deletion mapping to reference genes corresponding to deleted genes in the RIL143 deletion line. Reads mapped to multiple locations were split between locations. Genes with less than ten reads in the wildtype were eliminated to minimize the effect of random fluctuations in calling a deleted gene. We normalized the number of reads using the total number of reads and the length of the gene in kb, and then calculated the ratio between the normalized reads/kb in the mutant relative to the wildtype.

### 2.4. Growing conditions

The three single α-gliadin deletion lines and the combined *Δgli-A2 Δgli-D2* deletion were compared to the non-mutagenized RIL143 in the UC Experimental Field Station in Davis, CA (38° 32” N, 121° 46” W) located in the Sacramento Valley during the 2020-2021 and 2021-2022 growing seasons. The UC-Central Red sister lines with and without the *Δgli-D2* deletion were evaluated in the same location during the 2021-2022 and 2022-2023 growing seasons. The soil at this location is a deep Yolo loam (fine-silty, mixed, superactive, nonacid, thermic Mollic Xerofluvent). Sowing was done in November at a density of 300 grains per square meter (3 million grains per hectare) and managed with common agronomic practices. This included irrigations as needed and the application of ammonium sulfate fertilizer at the rate of 250 kg of N per hectare throughout the growing season. Due to moderate stripe rust susceptibility of the RIL143 line, the field experiments were treated with foliar fungicides during the growing season.

### 2.5. Breadmaking quality evaluations

We analyzed wheat quality at the California Wheat Commission Milling and Baking Laboratory (http://californiawheat.org/) using methods approved by the American Association of Cereal Chemist International (AACCI, Approved methods of analysis, 11th ed. AACC Intl., St. Paul http://methods.aaccnet.org/). Measured traits included test weight (AACCI 55-10.1) and grain and flour protein (AACCI 46-30.01). Before flour milling, 700 g of grain was tempered to 14% moisture and rested for 16-24 hours. Milling extraction rates were calculated against total products and reported on an “as is” moisture basis. Gluten strength was evaluated using both farinograph (AACCI 54-22.01) and mixograph (AACCI 54-40.02) assays. Finally, full baking tests were completed for all the samples (AACCI 10-12.01).

### 2.6. Proteomic studies

We extracted wheat proteins from 500 mg of Chinese Spring flour using 10 ml of 50% 1-propanol. Samples were manually inverted every 10 min over the course of 1h and centrifuged at 12,000 g for 10 min. The supernatant, which includes the alcohol-soluble protein fraction, was lyophilized and ground to a fine powder. The precipitate was subjected to several consecutive washing steps to minimize the presence of proteins in the glutenin fraction that were not covalently bound to the polymer. The washing steps included two 1-h washes with 50% 1-propanol at room temperature plus one wash at 60 °C for 1h. A final wash was performed with 2% SDS solution at 60 °C. Each washing step was followed by centrifugation and elimination of the supernatant. The final pellet was lyophilized and ground to a fine powder. Samples were then transferred to the UC Davis Genomic Center.

For each sample, 100 mg of wheat protein powder was solubilized into 5000 μL of solubilization buffer (5% SDS, 50 mM triethyl ammonium bicarbonate). This was placed inside tubes containing ceramic beads and homogenized with a MagNA Lyser instrument (Roche). The mix was centrifuges at 16,000 g and the supernatant containing the solubilized proteins was saved. Volumes containing 150 μg total protein from each sample were reduced with dithiothreitol and alkylated with iodoacetamide in 50mM triethylammonium bicarbonate (TEAB) buffer, then were subjected to a 2-enzyme digestion via suspension-trap (S-Trap) devices (ProtiFi). The enzymatic digestion included first the addition of trypsin 1:100 enzyme: protein (wt/wt) for 4 h at 37 °C, followed by additional trypsin digestion using the same wt/wt ratios for overnight digestion at 37 °C. Samples were then subjected to digestion with chymotrypsin, adding the same wt/wt ratio as for the tryptic digest, for 6 h at 37 °C. Peptides were eluted from S-Trap by sequential elution buffers of 100 mM TEAB, 0.5% formic acid, and 50% acetonitrile 0.1% formic acid. The eluted peptides were dried in a vacuum centrifuge and re-constituted in 0.1% trifluoroacetic acid.

Peptides were resolved on a Thermo Scientific Dionex UltiMate 3000 RSLC system using a PepSep analytical column (PepSep, Denmark): 150 μm x 8 cm C18 column with 1.5 μm particle size (100 Å pores), preceded by a PepSep C18 guard column, and heated to 40 °C. Each injection included 0.6 μg of total peptide. Separation was performed in a total run time of 60 min with a flow rate of 500 μL/min, and mobile phases including: A) water/0.1% formic acid and B) 80% ACN/0.1% formic acid. Gradient elution was performed from 60 min = 4% to 10% B over 4 min, from 10% to 46% B over 44 min, 46% to 99% B in 1.5 min, down to 4% B in 0.5 min followed by equilibration for 5 min.

Peptides were analyzed on an Orbitrap Exploris 480 instrument (Thermo Fisher Scientific, Bremen, Germany). Spray voltage was set to 1.8 kV, funnel radio frequency level at 45, and heated capillary temperature at 275 °C. The full MS resolution was set to 60,000 at m/z 200 and full MS automatic gain control (AGC) target was 300% with the injection time set to Auto. Mass range was set to 350–1500. For fragmentation spectra, isolation width was set at 1.6 m/z and normalized collision energy was set at 30%. The AGC target value was set to Standard with max injection time of 40 msec and we did TopN = 30 scans.

Mass spectrometry raw files were processed with FragPipe, v.21.1 (Nesvizhskii, A. I. lab, University of Michigan). For all searches, we used a protein sequence database including all manually curated gene families from cultivar Chinese Spring (iwgsc_refseqv1.1_manually_curated_gene_families.zip) at https://urgi.versailles.inra.fr/download/iwgsc/IWGSC_RefSeq_Annotations/v1.1/ and the Uniprot FASTA, CRAp, of common lab contaminants. Precursor mass tolerance was set to ± 20 ppm and 25 ppm for fragments. Trypsin was specified as protease. A maximum of two missing cleavages were allowed, the required minimum peptide sequence length was 4 amino acids, and the peptide mass was between 300-5000 Da. Carbamidomethylation on cysteine (+57.021 Da) was set as static modification, and oxidation on methionine was set as dynamic modification (+15.995 Da). To determine and control the number of false-positive identifications, MsFragger applies a target-decoy search strategy, using the concept of posterior error probability (PEP) to integrate multiple peptide properties, such as length, charge, and number of modifications into a single quantity reflecting the quality of a peptide spectral match. The reverse sequence library was used to control the false discovery rate (FDR) at less than 1% for peptide spectrum matches and protein group identifications. Decoy database hits, proteins identified as potential contaminants, and proteins identified exclusively by one site modification were excluded from further analysis. Label-free protein quantification was performed with the IonQuant algorithm v.10, but *Match Between Runs* feature (MBR) was disabled since each raw was submitted individually to Fragpipe search. We accepted identifications with at least one unique peptide. Among the identified peptides, we focused on diagnostic peptides for the α-gliadins with either seven (7-CYS) or six cysteines (6-CYS).

### 2.7. Statistical analyses

Yield and quality analysis assays for RIL143 and UC-Central Red were analyzed using least square means with years and blocks as random factors and blocks nested within years. Normality of the residuals was checked using the Shapiro-Wilk test. When the hypothesis of normality of residuals was rejected, data was transformed using different power transformations to restore normality. Adjusted means for the different traits obtained from the mutants were compared with the wildtype sister lines using a Dunnett multiple comparison test. The field experiments including the RIL143 control and four deletion lines were grown for two years in a randomized complete block design (RCBD) with five replications, whereas those including the UC-Central Red sister lines with and without the *Δgli-D2* deletion were grown for two years in a complete randomized design (CRD) with six replications.

## 3. Results

### 3.1. Haplotype analysis of the *Gli2* region

To determine the closest sequenced wheat genomes to the *GLI-A2*, *GLI-B2*, and *GLI-D2* loci in RIL143, we performed haplotype analyses for each region using SNPs detected in the α-gliadin genes and ten non-gliadin genes at each side of the *GLI2* loci. We generated exome capture data for RIL143 and compared it with exome capture data for 55 USA accessions obtained from the T3/Wheat database (WheatCAP project) and from 20 sequenced wheat genomes (Athiyannan et al. 2022; Avni et al. 2017; Maccaferri et al. 2019; Sato et al. 2021; Walkowiak et al. 2020).

The haplotype analysis for the 6A region (Fig. S1) revealed four haplotypes (HA1 to HA4), with HA1 and HA2 closer to each other and HA3 and HA4 clustered together (Data S2). The *GLI-A2* region from RIL143 was classified as haplotype HA1, which was also detected in the genomes of ArinaLrFor, Claire, Norin61, Robigus, and SY_Mattis. Interestingly, the *GLI-A2* loci from the available tetraploid and spelt wheats clustered together in a separate haplotype (HA2).

The haplotype analysis for the 6B region (Fig. S2) revealed eight haplotypes (HB1 to HB8) clustered in two major groups. The first one included haplotypes HB4-HB6 and the second one the other 5 haplotypes (Data S3). The 6B region from RIL143 was included in the largest HB1 haplotype, which was closest to the sequenced genomes of Kariega, Lancer and Mace.

Finally, the haplotype analysis for the 6D region (Fig. S3), which includes SNPs from six sequenced *Ae. Tauschii* genomes, revealed six haplotypes (HD1 to HD6). The *GLI-D2* region from RIL143 was included in haplotype H5, which was closer to the *Ae. tauschii* sequenced genomes than to the hexaploid wheat sequenced genomes, particularly in the region distal to the *GLI-D2* locus (Data S4). This result suggests that RIL143 6D region is either derived from a synthetic wheat, or that the HD2 haplotype is not present among the sequenced genomes by chance. Note that the *Gli-D2* region in CS RefSeq v1.0 and 2.0 is within the unassigned chromosome (Data S5).

Wheat cultivar Fielder is included in a different haplotype (HD2) than RIL143 based on the large number of SNPs in the region distal to *GLI-D2*, but is more similar to RIL143 in the α-gliadin sequences and the genes proximal to *GLI-D2* (Data S4). Six of the eight complete GLI-D2 proteins identified in Fielder were identical to their orthologs in RIL143 and the other two showed only one non-synonymous SNP each (Q206H in TraesFLD6D01G062300 and S43P in TraesFLD6D01G062600). Because of this sequence similarity, we selected the Fielder genome v1.0 as a reference for the *GLI-D2* gliadins in RIL143.

### 3.2. Distribution of epitopes and cysteines in the alpha gliadin loci

Another reason for the selection of the Fielder genome as a reference was its better assembly of the *GLI2* loci. All three *GLI2* loci seem to be complete, with no α-gliadins in the unassigned chromosomes. This better assembly likely reflects the improved technologies used to assemble the Fielder genome, which included long reads obtained with PacBio circular consensus sequencing using the HiFi approach (Sato et al. 2021). We manually curated the α-gliadins from Fielder, and the corrections and comments are provided in Data S5. After curation, we identified eight complete gliadins in *GLI-A2*, fourteen in *GLI-B2* and eight in *GLI-D2*.

We then characterized the number and location of the published lists of immunogenic nanomers (Chlubnova et al. 2023; Sollid et al. 2012; Sollid et al. 2020) and larger immunogenic peptides (Anderson et al. 2000; Juhasz et al. 2018; Shan et al. 2002) in each of the complete GLI2 α-gliadin proteins in Fielder (Fig. 1). These nanomers include the immunodominant peptides DQ2.5-glia-α1a (PFPQPQLPY), DQ2.5-glia-α1b (PYPQPQLPY) and DQ2.5-glia-α2 (PQPQLPYPQ) that produce consistently high responses in the majority of CeD patients (Jabri and Sollid 2017). Of the 105 immunogenic epitopes detected in Fielder α-gliadins, 53 (50.4%) are present in the *GLI-D2* locus (Data S5). The proportion is even larger among the immunodominant peptides, where 25 out of the 31 immunodominant epitopes (80.6%) are present in *GLI-D2*, 20% in *GLI-A2* and none in *GLI-B2* (Data S5).

**Figure 1.**
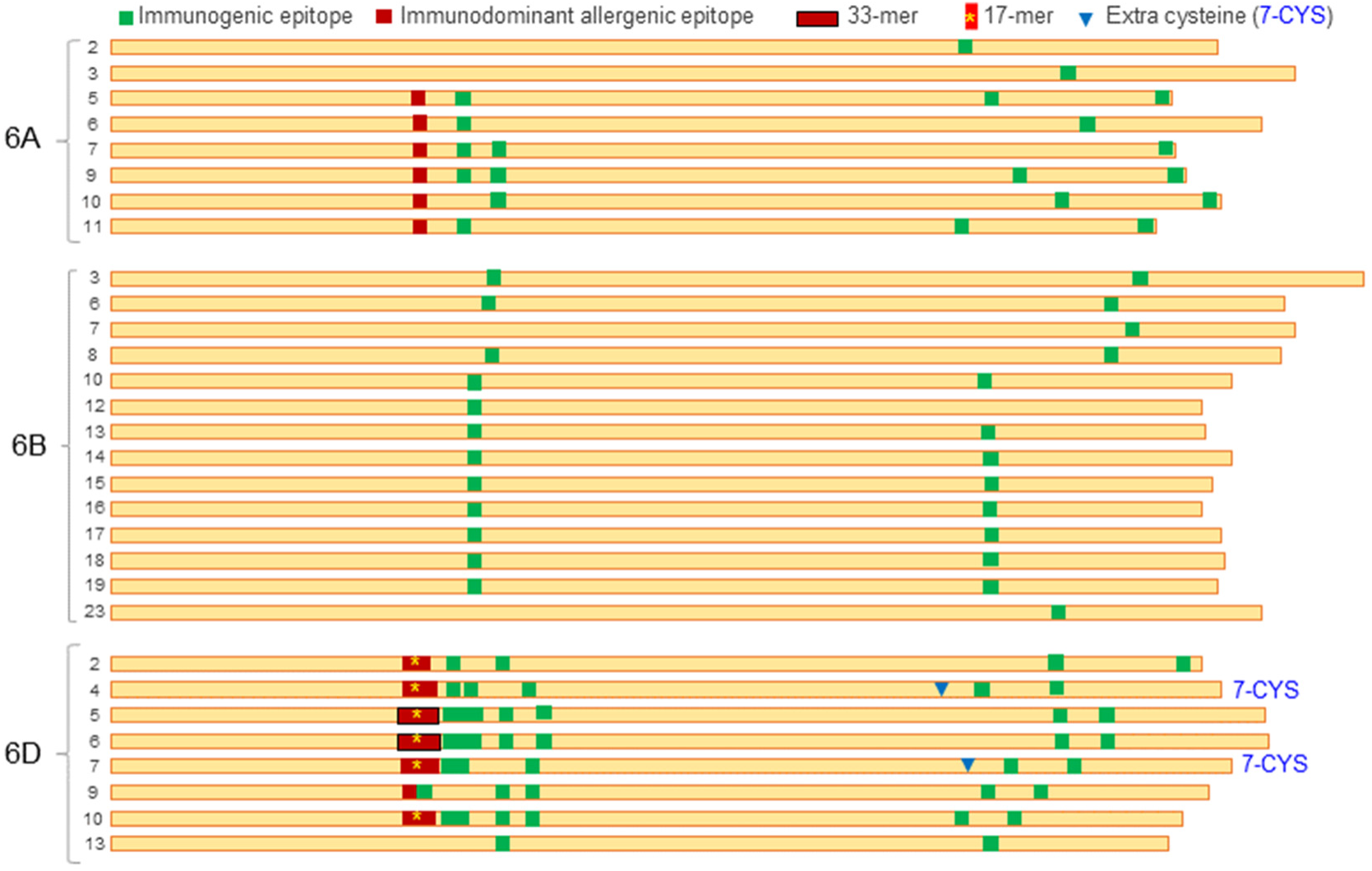
Complete alpha gliadin proteins from Fielder. All proteins showed 6 cysteines except for 6D-4 and 6D-7 (full gene names in Data S5), which have an additional cysteine indicated by a blue triangle. Squares represent CeD immunogenic epitopes, with those in red representing immunodominant epitopes and those in green less frequent epitopes (Chlubnova et al. 2023; Sollid et al. 2012; Sollid et al. 2020). Black rectangles indicate the immunogenic 33-mer including 5 overlapping immunodominant epitopes (Shan et al. 2002) and the * indicates the presence of the 17-mer within the immunodominant epitopes (Anderson et al. 2000).

In addition to these immunogenic nanomers, two of the α-gliadins in Fielder *GLI-D2* proteins (TraesFLD6D01G062600 and TraesFLD6D01G062700) contain a previously published 33-mer epitope that was shown to be highly immunogenic (Shan et al. 2002). This 33-mer includes five overlapping immunodominant nanomers and a 17-mer immunodominant peptide (QLQPFPQPQLPYPQPQ(S/P/L) (Anderson et al. 2000), which is also present in six out of the eight complete GLI-D2 proteins in Fielder (Fig. 1, Data S5). These 17-mer and 33-mer immunogenic peptides are conserved in the orthologous GLI-D2 proteins of RIL143, Chinese Spring, ArinaLrFor and *Ae. tauschii*, but were not detected in the *GLI-A2* or *GLI-B2* loci. Taken together, these results indicate that *Gli-D2* is the most immunogenic of the three α-gliadin loci in hexaploid wheat.

In addition to their higher immunogenicity, the α-gliadins from the *GLI-D2* locus from Fielder differ from those in the *GLI-A2* and *GLI-B3* loci by the presence of two α-gliadins with an unusual number of cysteines (CYS). The number of CYS is important because it determines the ability of these proteins to form intra- or inter-protein disulfide bonds, with even numbers favoring intra-chain bonds and odd numbers favoring both intra- and inter-chain bonds (Masci et al. 2002). All the complete α-gliadins found in the *GLI-A2* and *GLI-B2* loci in Fielder, ArinaLrFor, and Chinese Spring have six CYS located at relatively conserved distances (C-28/30-CC-11-C-76/119-C-7-C, Data S5). However, two to three of the GLI-D2 gliadins in these genomes have a seventh cysteine between CYS-4 and CYS-5 (Fig. 1, Data S5). Orthologous 7-CYS gliadins are also present in *Ae. tauschii GLI-D2* locus, indicating that these unusual gliadins were contributed by the diploid donor of the D genome. Based on their similarity with the *Ae. tauschii* and Fielder sequences, we were able to assign the 7-CYS α-gliadins present in the unassigned chromosome in Chinese Spring and ArinaLrFor to the *GLI-D2* locus. Using D genome specific primers and the *Δgli-D2* deletion, we also confirmed that the 7-CYS α-gliadins from RIL143 are present in the 6D chromosome.

### 3.3. Determination of the size of the radiation mutants with *GLI2* deletions

The PCR screening of the RIL143 radiation mutants with *GLI2* genome specific primers (Data S1) yielded one line with a deletion in *GLI-A2* (*Δgli-A2*), three with deletions in *GLI-B2* (*Δgli-B2*) and three with deletions in *GLI-D2* (*Δgli-D2*, Fig. 2). Polyacrylamide gel electrophoresis of the alcohol-soluble grain proteins from these lines showed the three *Δgli-B2* deletions shared the same missing bands, and the same was true for the three *Δgli-D2* deletions (Fig. 2). We selected one line for each genome and estimated the size of the deletions using exome capture.

**Fig. 2.**
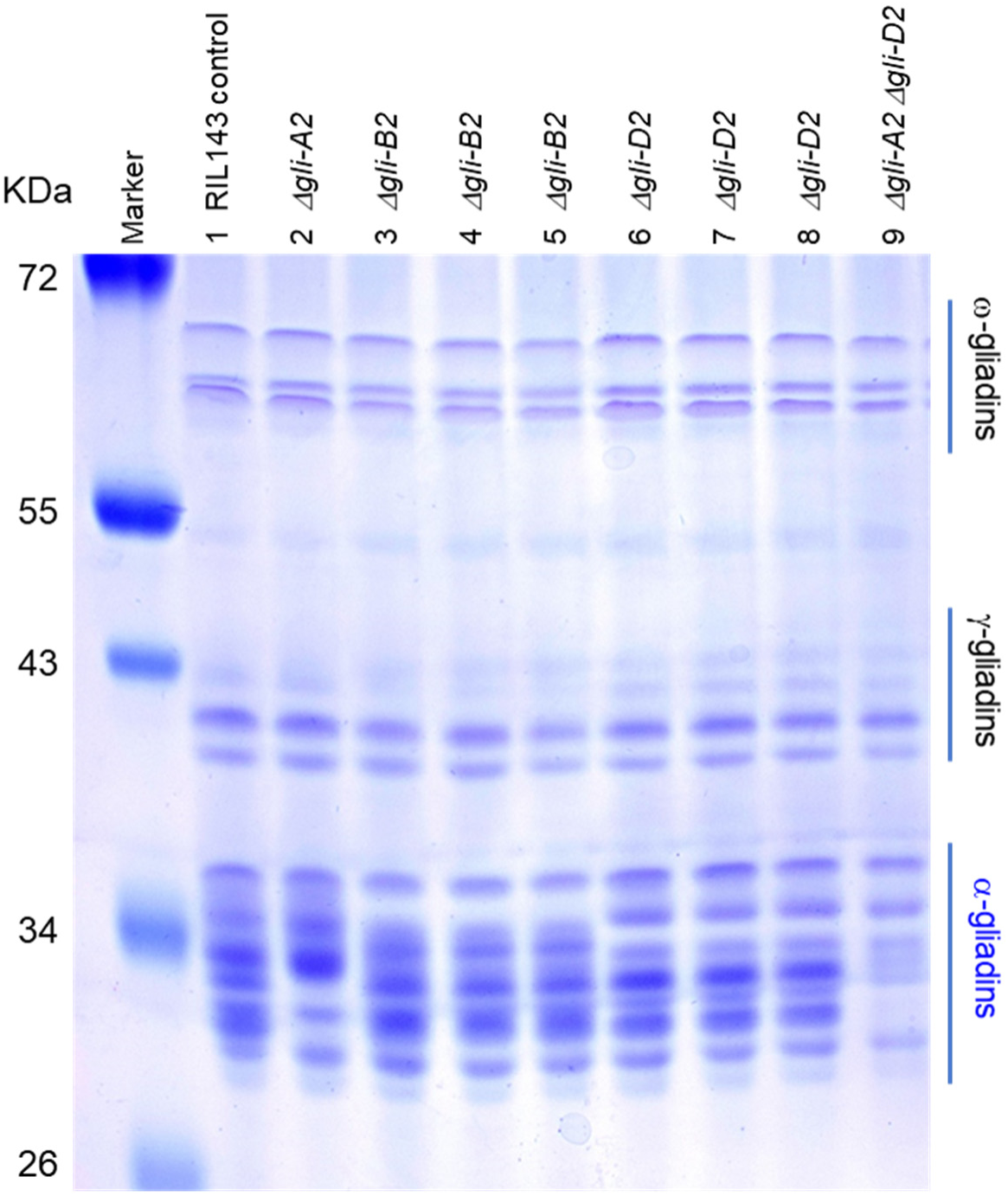
Polyacrylamide gel showing soluble proteins extracted with 50% propanol. Note the different α-gliadins missing in the *GLI-A2* (6A), *GLI-B2* (6B) and *GLI-D2* (6D) deletions. Some homoeologous bands overlap with each other and their deletion is only evident in the line carrying the 6A and 6D deletions (lane 9) or in changes in relative band intensities.

We used the coordinates of the Fielder genes flanking the borders of the deletions to estimate the minimum and maximum possible sizes of the deletions and the number of deleted gliadins and flanking genes (Table 1 and Fig. 3). We mapped the exome capture reads to the Fielder reference at high stringency and eliminated genes with less than 10 reads to minimize the effect of random fluctuations. We normalized the read count using the total number of reads and gene size (see Materials and Methods) and calculated the ratio between mutant and WT normalized reads. This ratio is expected to be close to 1 outside the deletion and close to 0 inside the deletion.

**Figure 3.**
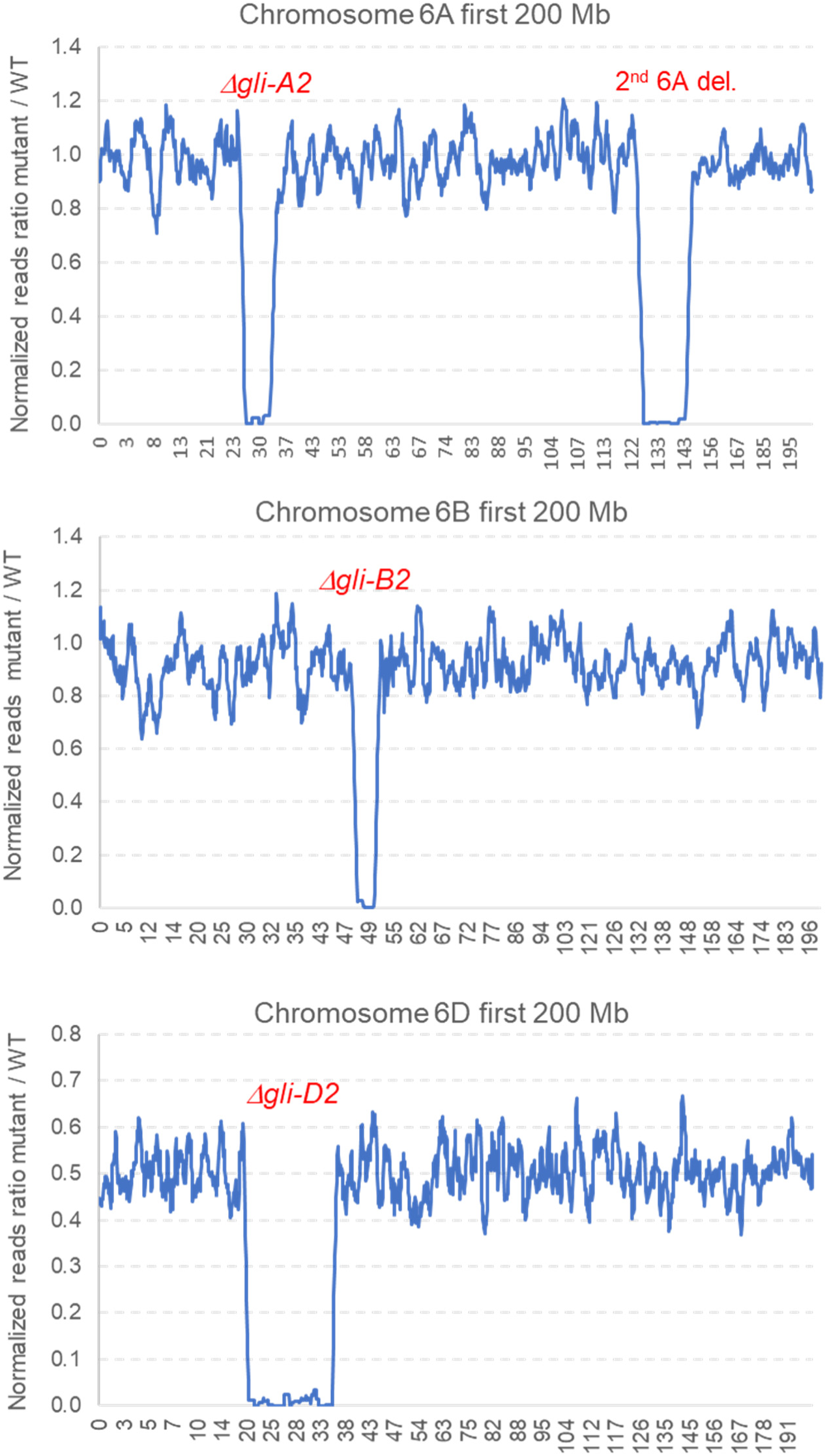
Graphical representation of the radiation deletions *Δgli-A2*, *Δgli-B2* and *Δgli-D2* using an overlapping moving window of ten genes (only first 200 Mb shown). A secondary deletion was detected on chromosome 6A. The values in the x-axis indicate the coordinates in Mb in the Fielder chromosomes.

**Table 1.** Characterization of radiation mutants that eliminate α-gliadins from chromosomes 6A, 6B and 6D.

| Fielder Coordinates | *Δgli-A2* | 6AS<br>2<sup>nd</sup> deletion | *Δgli-B2* | *Δgli-D2* |
| --- | --- | --- | --- | --- |
| Distal | 26,602,986 | 125,796,709 | 47,591,785 | 17,837,748 |
| Border | 27,500,174 | 127,291,438 | 47,799,566 | 19,674,383 |
| Proximal | 31,661,530 | 147,364,040 | 49,980,150 | 34,361,047 |
| Border | 31,710,947 | 148,295,698 | 50,229,076 | 34,738,339 |
| Size del. in Mb | 4.16-5.11 | 20.07-22.50 | 2.18-2.63 | 14.69-16.90 |
| *Gli2* excluding pseudogenes | 8 | 0 | 14 | 8 |
| Total non-*Gli2* genes | 79 | 93 | 24 | 262 |
| Gliadins outside deletion | 0 | NA | 1 | 0 |

Using this method, we estimated that the *Δgli-A2* deletion is between 4.2 to 5.1 Mb long (Table 1, Fig. 3, Data S6). This deletion includes all eight complete gliadins in *GLI-A2* and 79 other high-confidence annotated genes. We also mapped the exome capture reads of the RIL143 6A deletion to the closest genome of ArinaLrFor (Fig. S1, Data S2), which has an expanded *GLI-A2* locus with 14 complete gliadins, and confirmed that all 6A α-gliadins in RIL143 were deleted.

The RIL143 line carrying the *Δgli-A2* deletion has a secondary deletion estimated to be 20.1 to 22.5 Mb, which is located approximately 95.6 Mb proximal to *Δgli-A2* based on the Fielder genome coordinates (Table 1, Fig. 3, Data S6). We developed primers to detect the secondary deletion in 6A (Gap2F1 and Gap2R1, Data S1) and used them to test recombination events between the two deletions. Using these primers, we found that the secondary deletion was no longer present in a Fielder line in which we backcrossed the *Δgli-A2* deletion for four generations. This result demonstrates that these two deletions can be separated by recombination.

The *Δgli-B2* deletion was estimated to be between 2.2 and 2.6 Mb based on the Fielder genome coordinates (Table 1, Fig. 3, Data S6), and includes 13 of the 14 complete *GLI-B2* gliadins. The only exception is *TraesFLD6B01G102300*, which encodes an α-gliadin located 17 Mb proximal to the major *GLI-B2* cluster (at 67.4 Mb, Table 1 and Data S6). We confirmed that the RIL143 gene orthologous to *TraesFLD6B01G102300* was still present in the *Δgli-B2* deletion using gene specific primers (Data S1) and Sanger sequencing. In addition to the 13 missing gliadins, the *Δgli-B2* deletion includes 24 other high-confidence annotated genes.

Finally, we estimated that the 6D deletion in RIL143 is between 14.7 to 16.9 Mb long. The eight complete gliadins in this locus are all included in the deletion as well as other 262 annotated genes (Table 1, Fig. 3, Data S6). We intercrossed the three different α-gliadin deletions, but only the *Δgli-A2 Δgli-D2* combined deletion was fertile (Fig. 2, lane 9). Combinations of the 6B deletion with any of the other two resulted in plants with no anthers and multiple pistils that yielded no seeds (Fig. S4), and no plants with all three deletions were recovered. The three RIL143 individual deletions have been deposited in GRIN-Global and are publicly available without restrictions under germplasm accession numbers PI 704906 (*Δgli-A2*), PI 704907 (*Δgli-B2*), and PI 704908 (*Δgli-D2*).

### 3.4. Yield and agronomic field performance of the deletion lines

We evaluated the effect of the *Δgli-A2*, *Δgli-B2*, *Δgli-D2,* and the combined *Δgli-A2 Δgli-D2* on agronomic performance in the experimental field at UC Davis relative to the wildtype RIL143. Figure 4 presents the adjusted least square means across two years for grain yield (kg/ha, Fig. 4A), thousand kernel weight (g, Fig. 4B), test weight (kg/hl, Fig. 4C) and percent grain protein content (at 12% moisture, Fig. 4D). Additional data is available for plant height and heading time in supplemental Data S7 (RIL143).

**Figure 4.**
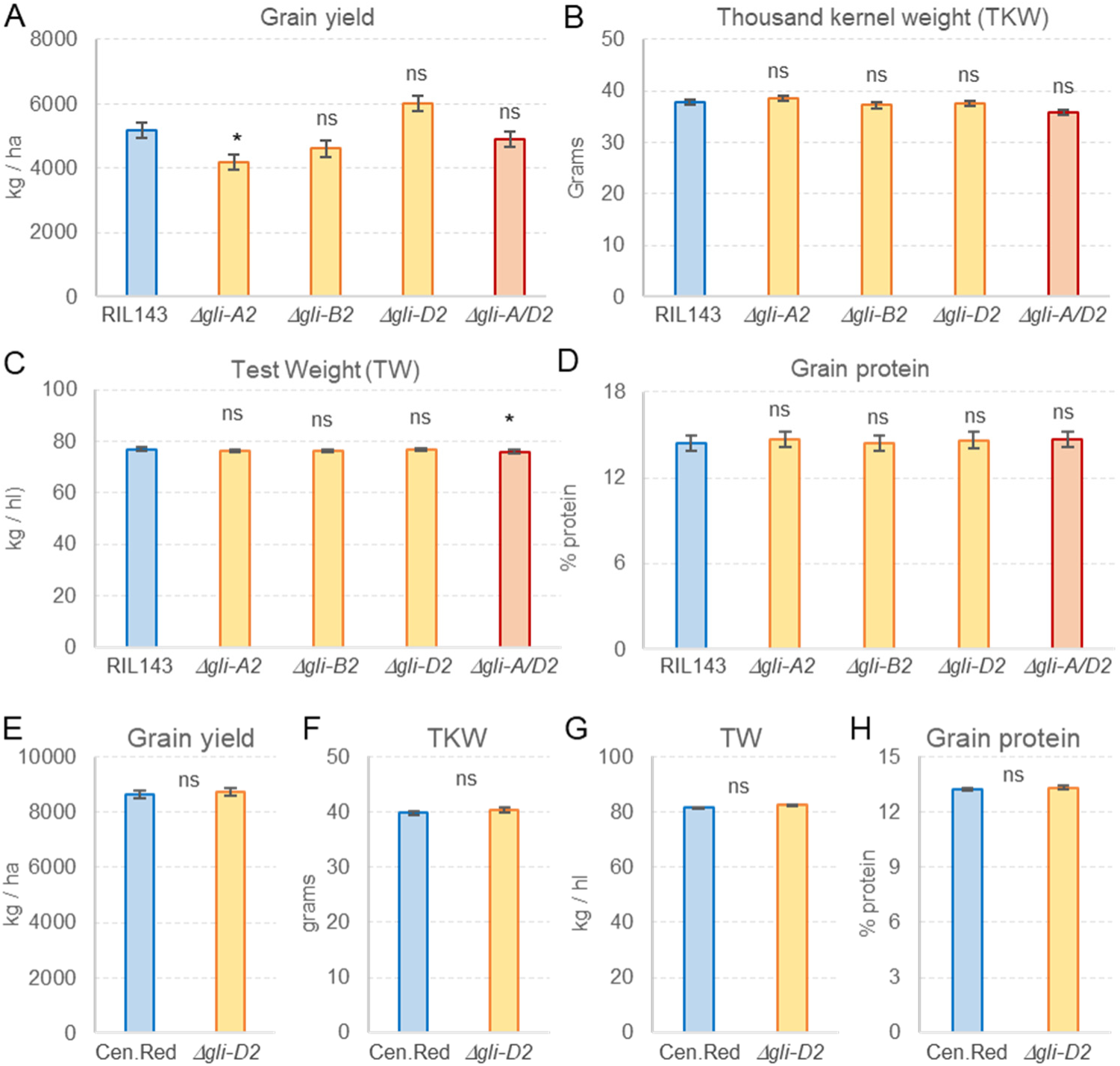
Field evaluation of RIL143 and UC-Central Red sister lines with and without *GLI2* deletions. (A-D) RIL143 and lines with deletions in chromosomes 6A (*Δgli-A2*), 6B (*Δgli-B2*), 6D (*Δgli-D2*), and both 6A and 6D (*Δgli-A/D2*) (n = 5). Adjusted means of the deletion lines were compared with the wildtype RIL143 using Dunnett tests. (E-H) UC-Central Red and its sister line with the *Δgli-D2* deletion (n = 6). Bars are least square means for the combined years and error bars are standard errors of the means (s.e.m.). *P* values are based on ANOVAs combining two years of field data, using years as blocks. Results ns= not significant, * = *P* <0.05. Raw data and statistical tests are available in Data S7 for A-D and Data S8 for E-H.

In general, the *GLI2* deletion lines showed non-significant differences from the wildtype RIL143 for all the traits described above (Fig. 4, and Data S7 and S8). One exception was an average 19% decrease in grain yield in the *Δgli-A2* deletion, which was marginally significant in the Dunnett test (*P* < 0.05, Fig. 4A). The *Δgli-D2* deletion showed slightly higher yields than the control but the differences were not significant. A slightly favorable effect of *Δgli-D2* was also observed in the combined *Δgli-A2 Δgli-D2*, which no longer showed the significant yield reduction detected in the *Δgli-A2* deletion alone (Fig. 4A). The combined *Δgli-A2 Δgli-D2* deletion also showed a significant reduction in plant height (Data S7), and a marginally significant effect on test weight (*P <* 0.05, Fig. 4C). However, the 1.2% increase in test weight is likely not agronomically relevant.

Since RIL143 is not a commercial variety and has limited yield potential and intermediate breadmaking quality, we introgressed the *Δgli-D2* deletion in the high-yielding and high-quality commercial line UC-Central Red by four backcrosses using the microsatellite marker BARC54 (Data S1). We initially focused on the *Δgli-D2* deletion, because it eliminates a significantly higher proportion of regular and immunodominant epitopes than the *Δgli-A2* and *Δgli-B2* deletions (Fig. 1). Additional data for plant height and heading time for UC-Central Red is available in Data S8. Similar to RIL143 deletion lines, the UC-Central Red sister lines with and without the *Δgli-D2* deletion show no significant differences in grain yield (Fig. 4E), thousand kernel weight (Fig. 4F), test weights (Fig. 4G), percent grain protein content (Fig. 4G), plant height or heading time (Data S8). Taken together, these data suggest that the introgression of the *Δgli-D2* deletion can be a useful tool for common wheat breeding programs.

### 3.5. Breadmaking quality evaluation of the radiation mutants

In contrast with the limited effects of the α-gliadin deletions on agronomic performance, we observed highly significant effects on breadmaking quality. In RIL143, these effects were larger in the *Δgli-D2* and combined *Δgli-A2 Δgli-D2* deletion, and smaller or not significant in the *Δgli-A2* and *Δgli-B2* deletions (Fig. 5A-D, Data S9). Relative to RIL143, *Δgli-D2* increased gluten index by 24% (Fig. 5A), farinograph stability by 129% (Fig. 5B), mixing peak time by 53% (Fig. 5C), and mixograph peak integral by 59% (Fig. 5D). The combined *Δgli-A2 Δgli-D2* deletion showed even larger increases (Fig. 5A-D).

**Figure 5.**
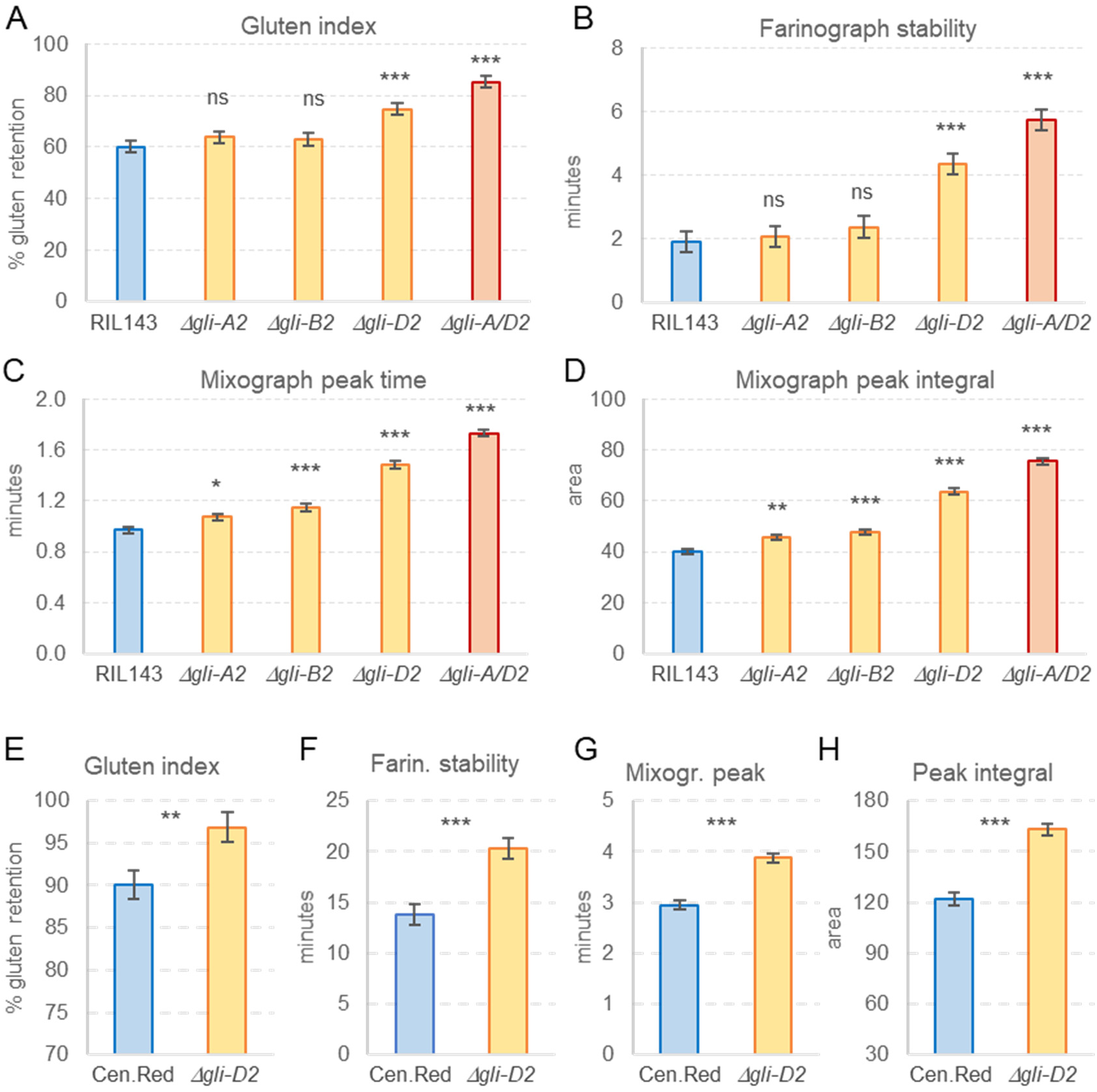
Breadmaking quality evaluation of RIL143 and UC-Central Red sister lines with and without *GLI2* deletions. (A-D) RIL143 and lines with deletions in chromosomes 6A (*Δgli-A2*), 6B (*Δgli-B2*), 6D (*Δgli-D2*), and both 6A and 6D (*Δgli-A2 Δgli-D2*) (n = 4). Adjusted means of the deletion lines were compared with the wildtype RIL143 using Dunnett tests. (E-H) UC-Central Red and its sister line with the *Δgli-D2* deletion (n = 6). Adjusted means were compared using an ANOVA with years as blocks. Bars are adjusted least-square means across years and error bars are standard errors of the adjusted means (s.e.m.). ns = not significant, * = *P* <0.05, ** = *P* <0.01, *** = *P* <0.001. Raw data and statistical tests are available in Data S9 for A-D and S10 for E-H.

In addition, the *Δgli-D2* and combined *Δgli-A2 Δgli-D2* deletion lines showed significant and positive effects on farinograph water absorption (4-5%), development time (62-72%) and mixing peak height (7%). Both deletion lines also exhibited beneficial effects in several baking tests including baking water absorption (4-5%), mixing time (16-31%), loaf volume (9-12%), dough handling (133-150%), crumb grain (49-62%) and bread symmetry (80-86%). These combined beneficial effects resulted in a significant increase of 59-65% in the overall baking score relative to RIL143 (Data S9). The only negative effect was a small 2-4% reduction in flour yield for the combined *Δgli-A2 Δgli-D2* deletion, which was not significant in *Δgli-D2* alone. This parameter was also marginally significant for *Δgli-A2* and *Δgli-B2* (*P* < 0.05).

The UC-Central Red line with the *Δgli-D2* deletion also showed positive and highly significant effects for gluten index (Fig. 5E), farinograph stability (Fig. 5F), mixing peak time (Fig. 5G), and mixograph peak integral (Fig. 5H, Data S10), confirming the beneficial effect of *Δgli-D2* on gluten strength. UC-Central Red has stronger gluten than RIL143, indicating that the *Δgli-D2* deletion can also benefit bread wheat lines with good breadmaking quality. However, as the baseline values for the gluten strength parameters in UC-Central Red are higher than those in RIL143 control, the percentage increases over the wildtype are relatively smaller in UC-Central Red than in RIL143.

The *Δgli-D2* deletion in UC-Central Red also showed additional beneficial effects in the sedimentation test (4.5%), bread mixing time (8.8%) and dough handling (13.3%). Loaf volume and baking scores were slightly higher in the *Δgli-D2* deletion than in the control, but the increases were not significant. The only negative effect was a marginally significant decrease (9%, *P* = 0.02) in total wet gluten (Data S10). In summary, the *Δgli-D2* deletion was associated with beneficial effects on gluten strength and breadmaking quality in the two genetic backgrounds with contrasting initial quality parameters.

### 3.6. *α*-gliadins with 7 cysteines are frequently incorporated in the gluten polymer

The previous results indicate that one or more genes located within the *Δgli-D2* deletion has a significantly negative impact on gluten strength in the wildtype, and that those genes should be different from their orthologs within the *Δgli-A2* and *Δgli-B2* deletions. Since only *Δgli-D2* includes 7-CYS α-gliadins, we hypothesized that those α-gliadins may contribute to the observed differences in gluten strength among deletion lines. This hypothesis predicts the incorporation of the 7-CYS α-gliadins into the gluten polymer by disulfide bonds.

To test this prediction, we separated proteins soluble in 50% 1-propanol from the gluten polymer by centrifugation, and thoroughly washed the gluten polymer to eliminate proteins that were not-covalently linked to the gluten polymer. We digested protein fractions with trypsin and chymotrypsin and analyzed the resulting fragments in an Orbitrap Exploris 480 instrument (Thermo Fisher Scientific, Bremen, Germany) (see Materials and Methods).

We aligned the 7-CYS and 6-CYS α-gliadins and identified several amino acid polymorphisms that were diagnostic of the 7-CYS (Fig. S5). We then focused on five peptides that included these diagnostic polymorphisms and that were present only in the 7-CYS α-gliadins (Fig. S6), and 19 control peptides that were present only in the 6-CYS α-gliadins detected in the D genome (Table 2).

**Table 2.**
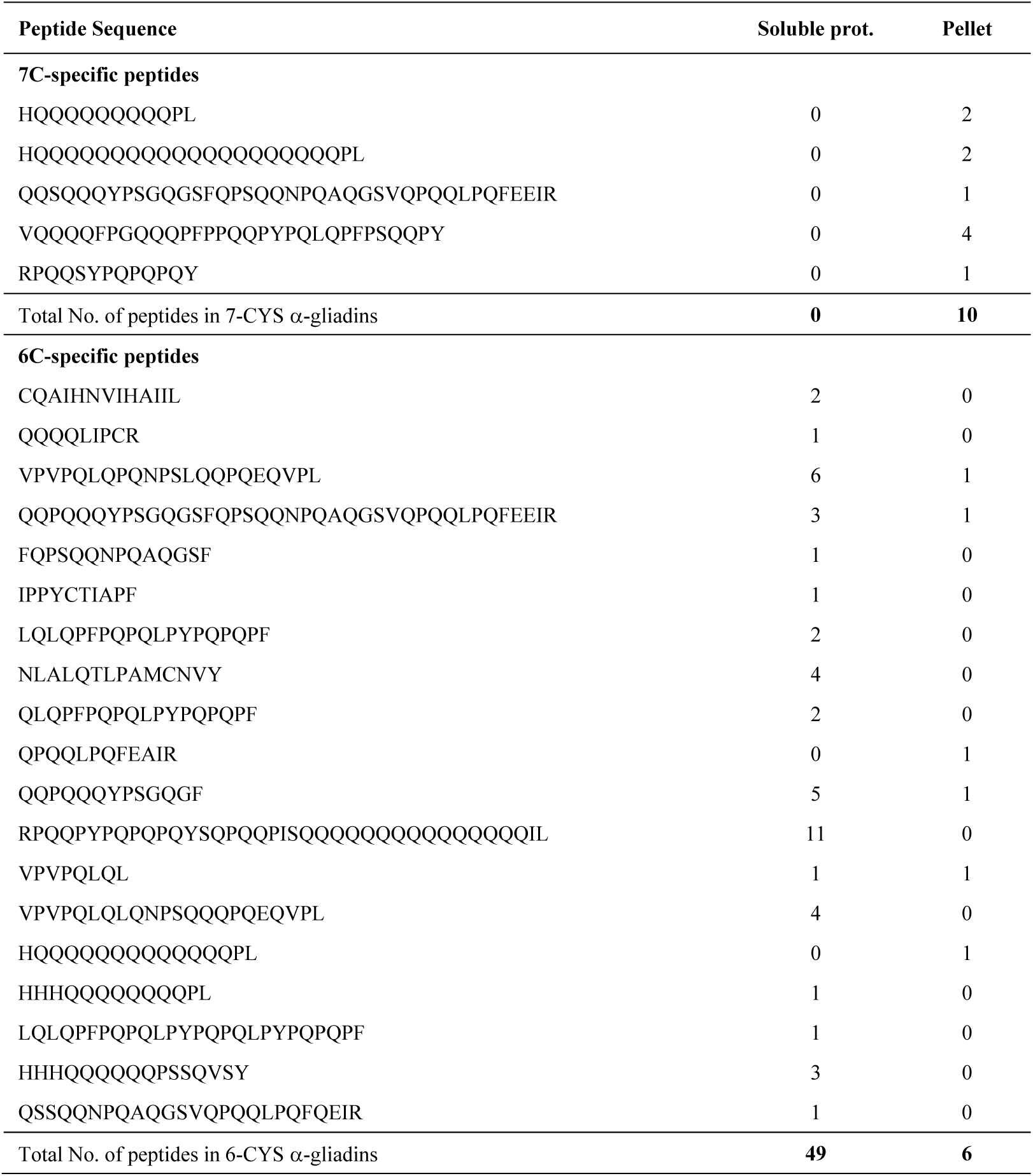
Frequency of *Gli-D2* digested peptides in the alcohol-soluble and insoluble fractions of gluten.

None of the 5 peptides mapped exclusively to the 7-CYS α-gliadins were detected in the alcohol-soluble fraction, and all 6 were present in the gluten fraction. A similar distribution was observed for 252 high molecular weight glutenin subunits (HMW-GS), which were detected mainly in the gluten fraction (96.8%). By contrast, 85.7% of the diagnostic 6-CYS peptides were detected in the alcohol-soluble fraction (Table 2). An Exact Fisher test confirmed that a highly significant departure (*P* < 0.0001) from a random distribution of the 7-CYS and 6-CYS peptides in the two fractions.

In summary, these results confirmed that the 7-CYS α-gliadins are incorporated into the gluten polymer more frequently than the 6-CYS α-gliadins, which are predominantly found in the alcohol-soluble phase.

## 4. Discussion

### 4.1. *α*-gliadins show extensive variation in copy number and polymorphism levels

The α-gliadins are present in wheat and closely related species of *Triticum* and *Aegilops*, but are not detected in rye or barley (Shewry et al. 1990), suggesting that they originated 3 to 5 million years ago (Gu et al. 2004). Comparisons of the α-gliadins in each of the three loci across the different wheat sequenced genomes revealed frequent variation in copy number, as expected for a tandemly repeated multigene family. For example, the *GLI-A2* locus in Fielder (HA3) has eight complete annotated α-gliadins, whereas fourteen are annotated in the same locus in ArinaLrFor (HA1). By contrast, the *GLI-B2* locus in Fielder (HB7) includes 14 complete α-gliadins, whereas only four are annotated in this locus in ArinaLrFor (HB4) (Data S5).

The *GLI-D2* locus showed reduced natural variation among the sequenced bread wheat genomes relative to the other two genomes (Data S4), which is consistent with the reduced variability of the *T. aestivum* D genome (Chao et al. 2007). We observed higher natural variation in *GLI-D2* among the *Ae. tauschii* sequenced genomes, and some of those polymorphisms were present in the exome captures of several hexaploid wheats, including RIL143 (Data S4). This result suggests that these varieties may carry *Ae. tauschii* introgression from the synthetic wheats developed at CIMMYT (Rosyara et al. 2019).

Fortunately, the field evaluations of the UC-Central Red sister lines with and without the *Δgli-D2* introgression indicate that this putative introgression is not associated with negative effects on grain yield, kernel weight, test weight or total grain protein content (Fig. 4). In summary, the haplotype analyses for the three *GLI2* loci presented in this study provide a useful framework to characterize the allelic diversity of the α-gliadins from tetraploid and hexaploid wheat.

### 4.2. *GLI-D2* includes *α*-gliadins with 7 cysteines that negatively affect gluten strength

An early sequencing study of wheat genomic clones including α-gliadins from cultivar Cheyenne confirmed that most α-gliadins encode proteins with six cysteines but also identified a few clones encoding α-gliadins with seven cysteines (Anderson et al. 1997). The authors of this study hypothesized that these unusual α-gliadins might be incorporated into the glutenin fraction. Using two-dimensional electrophoretic separation (A-PAGE vs SDS-PAGE), Masci et al. (2002) purified a glutenin fraction enriched in C-type low molecular weight glutenin subunits (LMW-GS) and mapped some of them to homeologous group 6. N-terminal amino acid sequencing revealed that 40% of the peptides from this fraction were similar to α-gliadins (Masci et al. 2002). Due to the limited protein sequence obtained by this method, this study was not able to determine the number of cysteines present in the gliadins incorporated into the gluten polymer.

We took advantage of the multiple sequenced wheat genomes available today to re-explore this question. We confirmed that, indeed, there are two to three α-gliadins encoding proteins with seven cysteines, and that these unusual gliadins are present exclusively in the D genome in all the examined bread wheat genomes. Moreover, we found these 7-CYS α-gliadins in the *Ae. tauschii* genomes confirming that they were contributed to bread wheat by the diploid donor of the D genome (Data S5). The six cysteines found in the canonical α-gliadins form three intra-molecular disulfide bonds (Muller et al. 1995), resulting in monomeric proteins with no cysteines available to form covalent bonds with the gluten polymer. It was previously hypothesized that 7-CYS α-gliadins could form one intermolecular disulfide bond with the gluten polymer and act as terminators of the growing glutenin polymer chains, resulting in shorter gluten macropolymers (Kasarda et al. 1987; Masci et al. 2002; Shewry and Lafiandra 2022). Since the proportion of gluten macropolymer with apparent size above 158 kDa is positively correlated with gluten strength (Gupta et al. 1993), the presence of 7-CYS α-gliadins is predicted to be detrimental to gluten strength and breadmaking quality. Our proteomics studies demonstrate that, indeed, the 7-CYS α-gliadins are preferentially incorporated to the gluten macropolymer (Table 2).

The effect of a *GLI-D2* deletion on gluten polymer size was previously determined in the cultivar Xiaoyan 81 (Li et al. 2018). This study found that the presence of the deletion was associated with a significant increase in the proportion of large glutenin macropolymers (>250 kDa) and unextractable proteins (Li et al. 2018). We re-analyzed the published sequences of the α-gliadins in Xiaoyan 81 (Li et al. 2018) and confirmed the presence of at least one α-gliadin with seven cysteines. We hypothesize that the deletion of this 7-CYS α-gliadin may have contributed to the increased polymer size observed in the *GLI-D2* deletion.

The *GLI-D2* deletion in Xiaoyan 81 was also associated with significant increases in dough development and stability times and with increases in loaf volume relative to the control line (Li et al. 2018). The authors hypothesized that these favorable changes in the deletion line were the result of increases in the proportion of glutenins relative to gliadins, likely caused by compensatory higher expression of the glutenin genes in the deletion line (Li et al. 2018). Although our *Δgli-D2* deletion also showed improved gluten strength, those large effects were not detected in *Δgli-A2* or *Δgli-B2* which also have a large number of deleted α-gliadins (Fig. 5, Data S9). These results suggest that the increase in gluten strength associated with the *Δgli-D2* deletion may not be just a compensatory increase in the expression of glutenin genes, but a more specific effect of the α-gliadins present in the D genome.

Taken together, the presence of 7-CYS α-gliadins only in the D genome, the stronger effect of the *Δgli-D2* deletion on gluten strength relative to *Δgli-A2* or *Δgli-B2*, the demonstration that the 7-CYS α-gliadin are preferentially incorporated into the gluten polymer, and the positive effect of the Xiaoyan 81 *GLI-D2* deletion on the proportion of large gluten macropolymer (Li et al. 2018), all support the hypothesis that the 7-CYS α-gliadins can act as chain extension terminators and have a negative impact on gluten strength and breadmaking quality.

### 4.3. A path to reduced immunogenicity wheat

Although 30 to 40% of the human population express one or both CeD predisposing HLA-DQ allotypes HLA-DQ2 and HLA-DQ8, the prevalence of CeD is close to 1%. This difference indicates that additional genetic and environmental factors are involved in the onset of the disease (Abadie et al. 2024). Several studies suggest that one of those environmental factors is the level of exposure to immunogenic epitopes (Ivarsson et al. 2000; Koning 2012; Mearin et al. 1983; Megiorni et al. 2009). Therefore, a reduction in the level of wheat immunogenic epitopes may reduce the incidence of CeD at a population level. A logical start for this reduction path is the elimination of the immunodominant epitopes that result in high immune responses in the majority of the CeD patients. The elimination of immunogenic proteins with limited impact on breadmaking quality is an additional useful criterion to accelerate the deployment of loci with reduced immunogenic proteins.

The α-gliadins are a desirable initial target since the *GLI2* loci segregate independently of the LMW-GS and HMW-GS, which have a major impact on gluten strength. In particular, the *GLI-D2* locus has on average 55% of the total epitopes and 81% of the immunodominant nanomers present in the α-gliadins (Fig. 1, Data S5, average across Fielder, Chinese Spring and ArinaLrFor). In addition, only the *GLI-D2* locus carries the highly immunogenic 17-mer (Anderson et al. 2000) and 33-mer peptides (Shan et al. 2002) formed by multiple overlapping immunodominant nanomers. The presence of the 33-mer peptide only in the *GLI-D2* locus was also reported in the Italian variety Pegaso (Camerlengo et al. 2017).

In this study, we describe a *Δgli-D2* deletion that eliminates all the α-gliadins from the *GLI-D2* locus and, therefore, all the 17-mer and 33-mer highly immunogenic epitopes and ~81% of the immunodominant nanomers present in the α-gliadins. Although this deletion includes 262 genes in addition to the α-gliadins, we observed no negative effects on grain yield, grain protein content, test weight or kernel weight in the two genetic backgrounds evaluated over two years. More importantly, the *Δgli-D2* deletion showed a positive impact on gluten strength and breadmaking quality that was consistent in both low- and high-quality genetic backgrounds across the two years of replicated trials. A similar positive effect on gluten strength and breadmaking quality was observed in an independent *Δgli-D2* deletion in the Chinese bread wheat variety Xiaoyan 81 across four environments, which was also validated in a different genetic background with stronger HMW-GS alleles (Li et al. 2018). Increases in dough development time and stability measured by micro-mixograph tests were also observed in greenhouse experiments comparing Chinese Spring with lines carrying large terminal deletions of the short arm of chromosome 6DS. However, variable results were observed for other traits, likely associated with the large size of these deletions, which eliminated more than half of the short arms of chromosomes 6A, 6B and 6D (van den Broeck et al. 2009).

Although immunogenic epitopes are present in the α-gliadins from all three *GLI2* loci, the immunodominant epitopes are restricted to the *GLI-D2* and *GLI-A2* loci (Fig. 1, Data S5). Therefore, the simultaneous introgression of the *Δgli-D2* and *Δgli-A2* deletions can be used to further reduce the levels of immunodominant epitopes in hexaploid wheat. In this study we show that the combination of these two deletions has no significant effects on grain yield (Fig. 4) and has the same beneficial effects on gluten strength and breadmaking quality as the single *Δgli-D2* deletion (Fig. 5). Different deletions for *GLI-A2* from cultivar ‘Raeder’ (Lafiandra et al. 1987) and for *GLI-D2* from cultivar ‘Saratovskaja 29’ (Redaelli et al. 1994) were combined in the Italian bread wheat variety Pegaso, but the size of these deletions and their effect on breadmaking quality was not evaluated (Camerlengo et al. 2017).

We were not able to combine the α-gliadin deletions in the three genomes because the combinations including *Δgli-B2* both showed abnormal flowers (Fig. S4) and were completely sterile. Wheat is a young polyploid species and often tolerates deletions of one and sometimes two homeologs, but deletions of three homeologs are expected to have the same effects as in a diploid species (Uauy et al. 2017). The risk of deleting all three homeologs of essential or important genes limits the use of large deletions to obtain triple null mutants for the α-gliadins. Smaller α-gliadin deletions or CRISPR editing of the α-gliadins in at least one of the three genomes will be likely required to generate simultaneous deletions of all α-gliadins and study their effect on quality and agronomic performance.

Although the combined *Δgli-A2 Δgli-D2* deletions would eliminate the known immunodominant epitopes present in the α-gliadins, there are additional immunodominant epitopes (DQ2.5-glia-ω1 and DQ2.5-glia-ω2) in the ω-gliadins in the *GLI-A1*, *GLI-B1*, and *GLI-D1* loci. Most of these immunodominant epitopes are present in the *GLI-D1* locus. Fortunately, there is a published 1D deletion including part of the *GLI-D1* locus that eliminates the ω-gliadins but not the LMW-GS, which was originally identified in the French cultivar ‘Darius’ (Branlard et al. 2003) and was then transferred to ‘Pegaso’ (Camerlengo et al. 2017). We are currently combining this deletion with *Δgli-A2* and *Δgli-D2* and evaluating its effect on grain yield and quality. There are other CRISPR-induced deletions and knockouts including multiple gliadin loci (Jouanin et al. 2019; Sánchez-León et al. 2018).

One useful resource is a CRISPR line in the cultivar Fielder that eliminated all the ω-gliadins and some γ-gliadin (Yu et al. 2024). These deletions were associated with a significant increase in the ratio of glutenins to gliadins, the ratio of polymeric to monomeric proteins, and the proportion of gluten macropolymer in total protein. These changes were associated with significant increases in mixograph peak time and farinograph stability time, two parameters associated with improved gluten strength (Yu et al. 2024). These results, together with the effects of the α-gliadin deletions reported in this study, indicate that a large proportion of the immunogenic α-, ω- and γ-gliadins can be eliminated without negatively affecting breadmaking quality and, in some cases, even generating positive effects in gluten strength and other breadmaking quality parameters.

Even if all the immunodominant epitopes are eliminated, the resulting plants will not be safe for people that already have CeD because of the presence of residual epitopes. The elimination of all epitopes will require the modification of LMW-GS and HMW-GS that are essential for gluten quality, and that will need to be replaced by transgenic functional versions without the epitopes to restore the lost breadmaking quality. Although the development of CeD-safe wheats will likely have to wait for a broader acceptance of transgenic technologies, the development of wheats lines with reduced immunogenicity is an achievable intermediate objective.

The development of wheat lines without immunodominant epitopes will be useful to quantify the impact of this reduction on the onset of the CeD disease in genetically predisposed individuals, and the value of these lines to reduce the incidence of CeD at the population level. Meanwhile, the *Δgli-A2* and *Δgli-D2* lines developed in this study can be combined to eliminate the immunodominant epitopes from the α-gliadins. In particular, the incorporation of the *Δgli-D2* deletion into commercial bread wheat varieties will simultaneously eliminate many immunodominant epitopes and improve gluten strength and breadmaking quality. To accelerate the deployment of these deletions in wheat breeding programs, we have deposited the RIL143 *Δgli-A2* (PI 704906), *Δgli-B2* (PI 704907), and *Δgli-D2* (PI 704908) deletions in GRIN-Global and made them publicly available without any restrictions.

## Abbreviations

AACCI: American Association of Cereal Chemists International
AGC: automatic gain control
CYS: cysteine
GRIN: Germplasm Resources Information Network
HMW-GS: High-molecular-weight glutenin subunits
LMW-GS: Low-molecular-weight glutenin subunits
MS: mass spectrometry
PAGE: polyacrylamide gel electrophoresis
RSLC: rapid separation liquid chromatography
SDS: sodium dodecyl sulfate
SNP: single nucleotide polymorphism
UC: University of California
WT: wildtype

## Acknowledgements

We thank Huiqiong Lin (UC Davis) for her help with the floret pictures of the Kronos wildtype and the combined deletion lines.

## Author Contributions

MGR-writing - original draft, data curation, investigation, visualization, statistical analyses formal analysis. WZ-investigation, methodology, writing, reviewing & editing. XZ-methodology. JZ-formal analysis, writing, reviewing & editing. GG-data curation, methodology, writing – reviewing & editing. GB-data curation, bioinformatics methodology, reviewing & editing. JH-field methodology, data curation, reviewing & editing. TV and CC-quality methodology, supervision, resources, reviewing & editing. JD-conceptualization, visualization, statistical analyses, funding acquisition, project administration, supervising, formal analysis, writing – reviewing & editing.

