## Supplementary Figures for "Deletion of wheat alpha-gliadins from chromosome 6D improves gluten strength and reduces immunodominant celiac disease epitopes"

**Fig. S1.** Haplotype analysis for the α-gliadin locus and flanking regions on chromosome 6A. Tetraploid accessions are indicated in blue. Sequenced genomes are indicated in bold. Data S2.

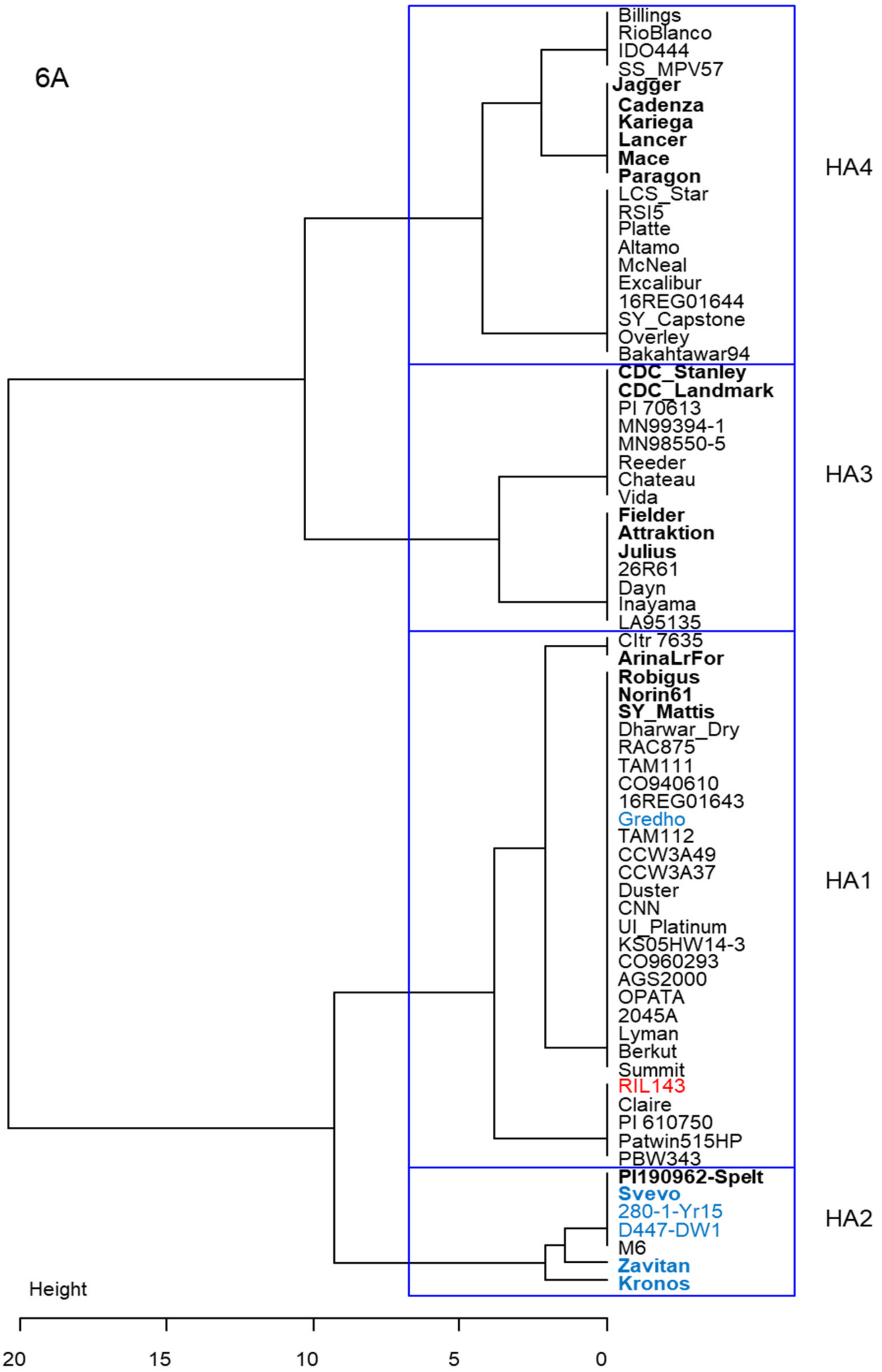

**Fig. S2.** Haplotype analysis for the α-gliadin locus and flanking regions on chromosome 6B. Tetraploid accessions are indicated in blue. Sequenced genomes are indicated in bold. Data S3.

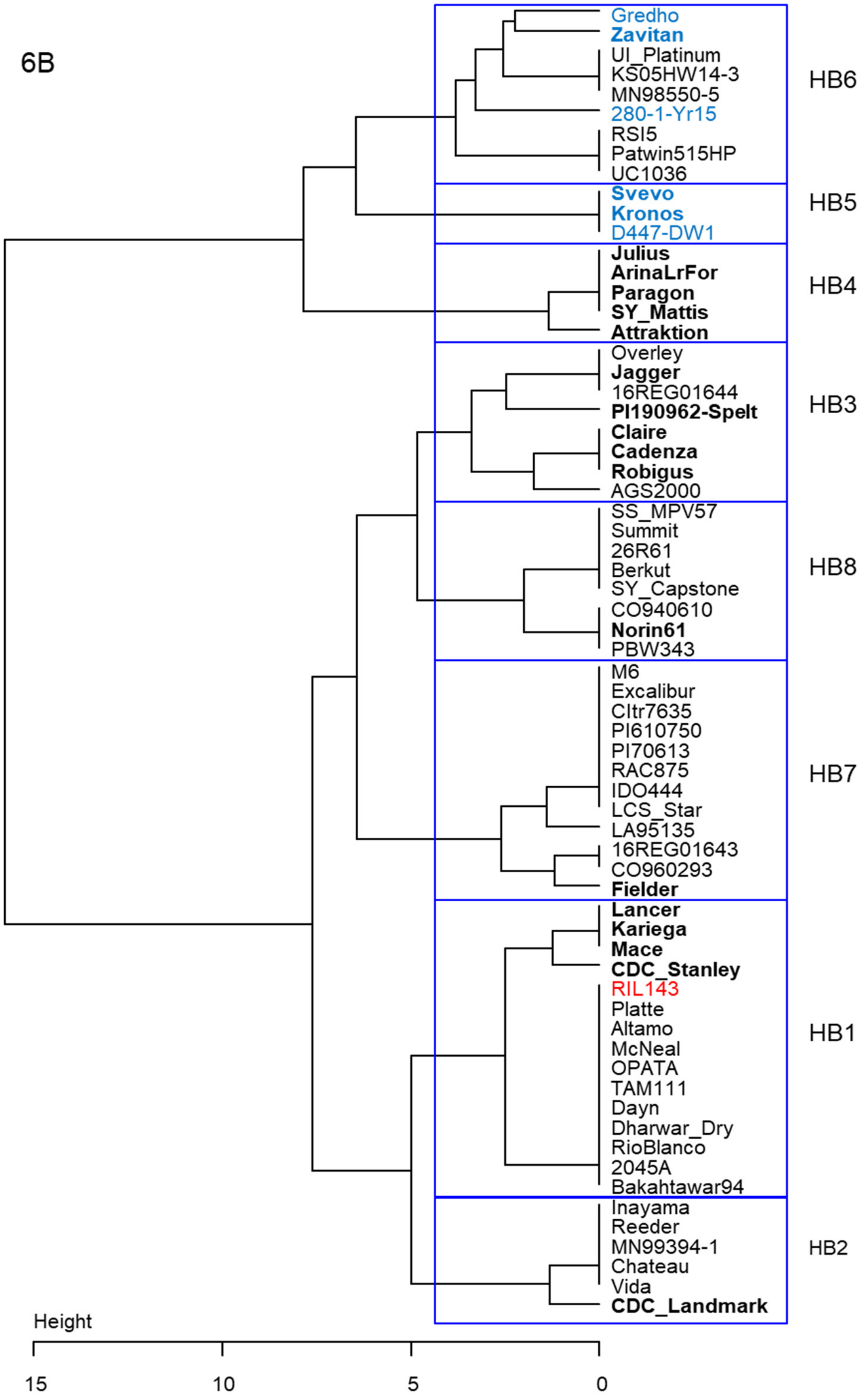

**Fig. S3.** Haplotype analysis for the α-gliadin locus and flanking regions on chromosome 6D. *Ae tauschii* accessions are indicated in green. Sequenced genomes are indicated in bold. Data S4.

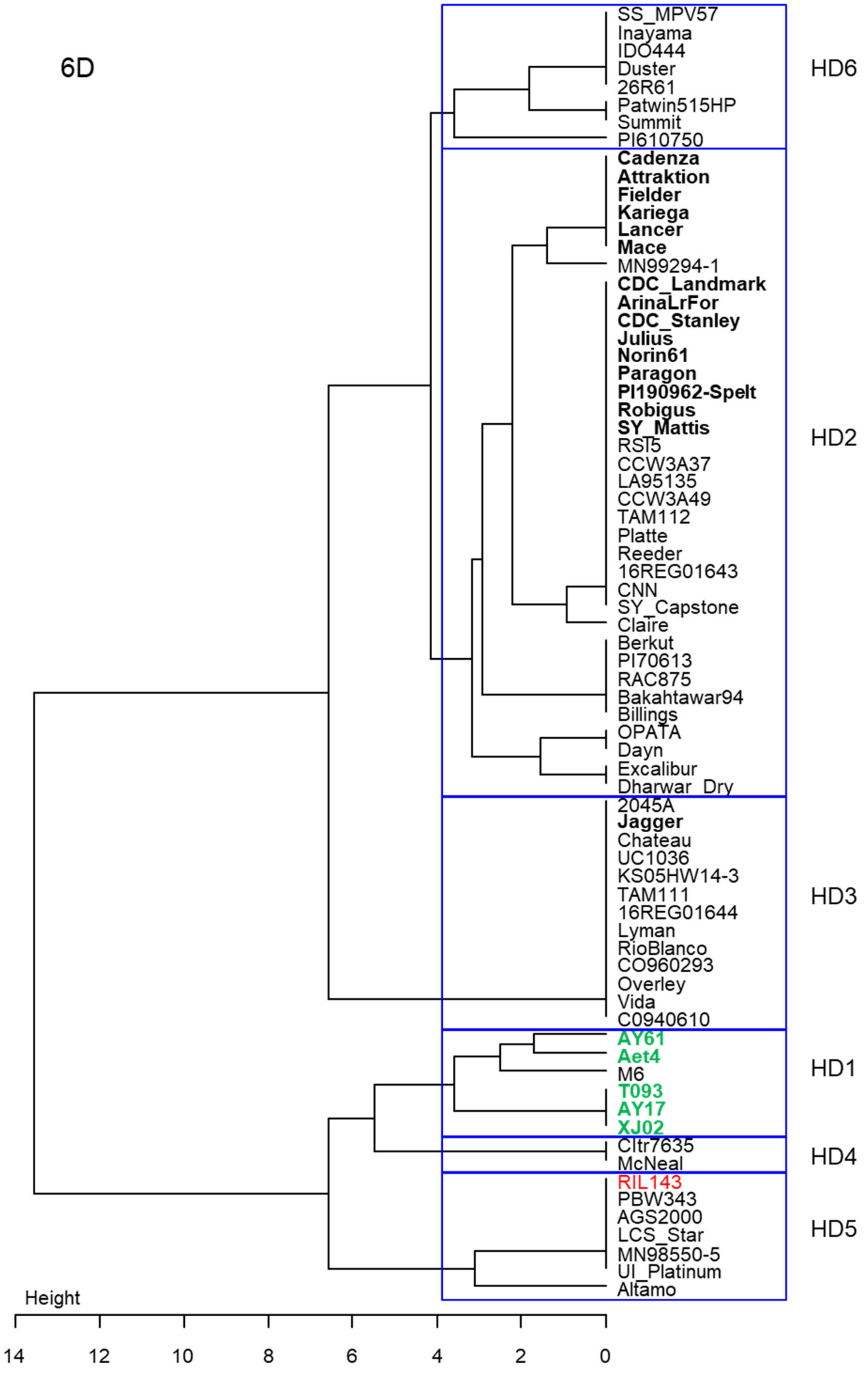

**Fig. S4.** Wildtype Kronos and *Δgli-B1 Δgli-D1* combined deletion line. The left panel shows normal floral organs in wildtype Kronos and the right panel shows abnormal sterile flowers with no stamens and multiple pistils in the combined *Δgli-B1 Δgli-D1* deletion line.

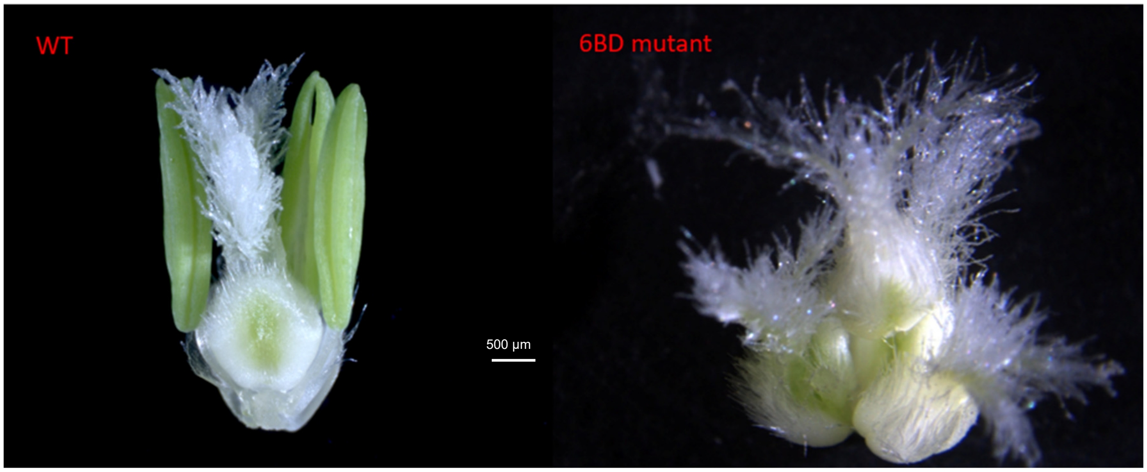

**Fig. S5.** Alignment between 7-CYS and 6-CYS α-gliadins from Chinese Spring detected in the proteomics study. Genes indicated as CSU were assigned to chromosome 6D based on high similarity to colinear genes in Fielder. Amino acids that separate the 7-CYS from the rest are marked in red and highlighted in yellow.

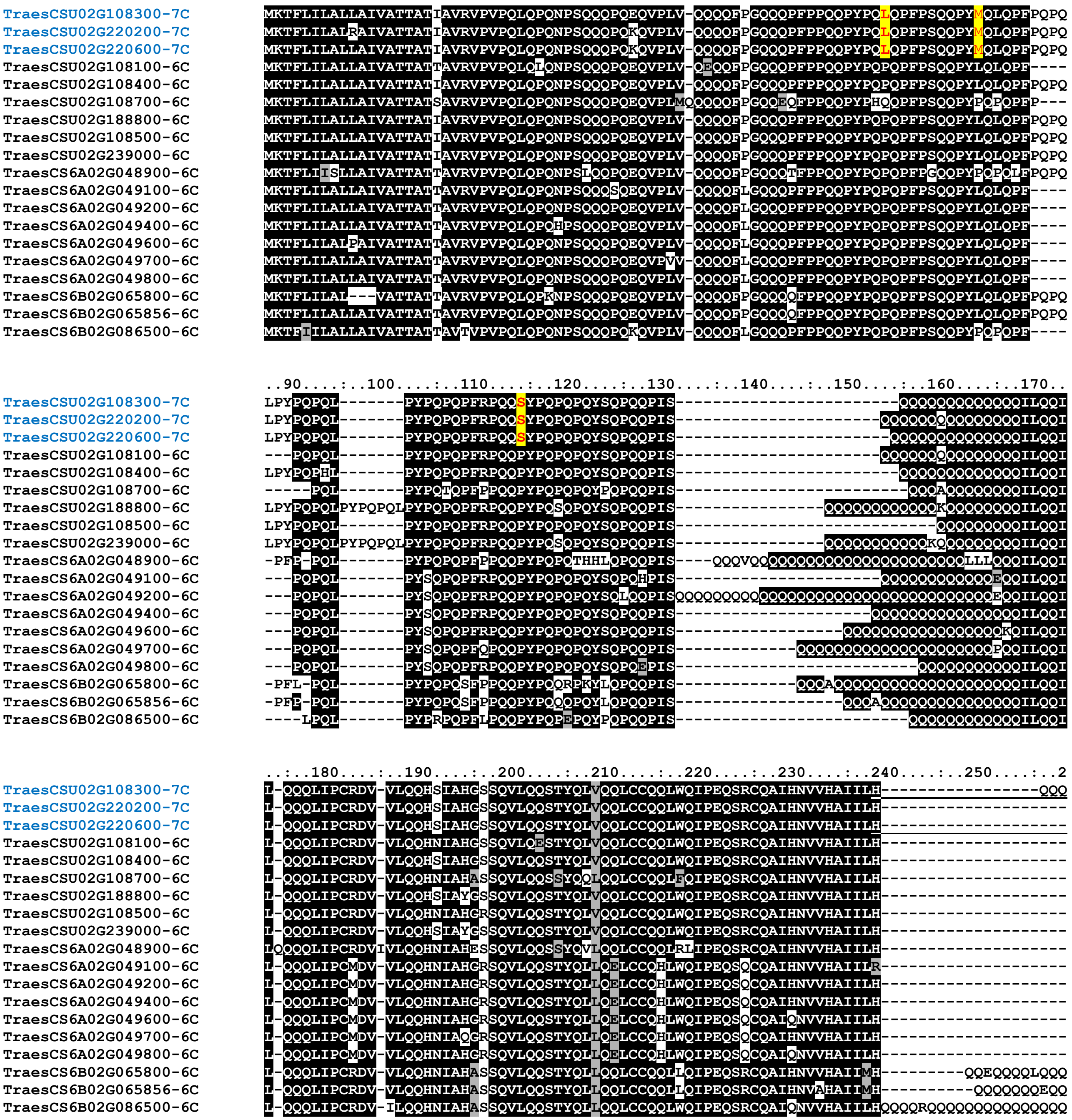

```
                          60....:..270....:..280....:..290....:..300....:..310....:..320....:..330....:..340....
TraesCSU02G108300-7C      QQQQQQQQQQQQQQQQQPLSQVCFQQSQQQYPSGQGSFQPSQQNPQAQGSVQPQQLPQFEEIRNLALETLPAMCNVYIPPYC--TI
TraesCSU02G220200-7C      --QQQQQQQQQQQQQQQPLSQVCFQQSQQQYPSGQGSFQPSQQNPQAQGSVQPQQLPQFEEIRNLALETLPAMCNVYIPPYC--TI
TraesCSU02G220600-7C      --------QQQQQQQQQPLSQVCFQQSQQQYPSGQGSFQPSQQNPQAQGSVQPQQLPQFEEIRNLALETLPAMCNVYIPPYC--TI
TraesCSU02G108100-6C      -QQHHHHQQQQQQQQQQPLSQVSFQQPQQQYPSGQGFFQPSQQNPQAQGSFQPQQLPQFEAIRNLALQTLPAMCNVYIPPYC--TI
TraesCSU02G108400-6C      ----QQQQQQQQQQQQQPLSQVSFQQPQQQYPSGQGSFQPSQQNPQAQGSVQPQQLPQFEEIRNLALETLPAMCNVYIPPYC--TI
TraesCSU02G108700-6C      ---------HHQQQQQQPSSQVSYQQPQEQYPSGQGSFQSSQQNPQAQGSVQPQQLPQFQEIRNLALQTLPAMCNVYIPPYCSTTI
TraesCSU02G188800-6C      -----QQQQQQQQQQQQPLSQVSFQQPQQQYPSGQGSFQPSQQNPQAQGSVQPQQLPQFEEIRNLALETLPAMCNVYIPPYC--TI
TraesCSU02G108500-6C      -------HHQQQQQQQQPLTQVSFQQPQQQYPSGQGSFQPSQQNPQAQGSVQPQQLPQFEEIRNLALETLPAMCNVYIPPYC--TI
TraesCSU02G239000-6C      ----QQQQQQQQQQQKQPLSQVSFQQPQQQYPSGQGSFQPSQQNPQAQGSVQPQQLPQFEEIRNLALETLPAMCNVYIPPYC--TI
TraesCS6A02G048900-6C     -QQQQQQQQQQQQGQHQPSSQVSYQQPQQQYPSGQGSFQPSQQNPQAQGFVQPQHLPQLEEISNLALQTLPAMCNVYIPPYCSTTI
TraesCS6A02G049100-6C     ---------QQQKQQQQPSSQVSFQQPQQQYPLGQGSFRPSQQNPQAQGSVQPQQLPQFEEIRNLALQTLPAMCNVYIPPYC--TI
TraesCS6A02G049200-6C     --------QQQKQQQQQPSSQVSFQQPQQQYPLGQGSFRPSQQNPQAQGSVQPQQLPQFEEIRNLALQTLPAMCNVYIPPYC--TI
TraesCS6A02G049400-6C     ---------QQQKPQQQPSSQVSFQQPLQQYPLGQGSFRPSQQNPQARGSVQPQQLPQFEEIRNLALQTLPAMCNVYIPPYC--TI
TraesCS6A02G049600-6C     ---------QQQKQQQQPSSQVSFQQPLQQYPLGQGSFRPSQQNPQDQGSVQPQQLPQFEEIRNLALQMLPAMCNVYIPPYC--TI
TraesCS6A02G049700-6C     ----QQQKQQQQQQQQQPSSQVSFQQPQQQYPLGQGSFRPSQQNPQAQGSVQPQQLPQFEEIRNLALQTLPAICNVYIPPYC--TI
TraesCS6A02G049800-6C     ---------QQQKQQQQPSSQVSFQQPLQQYPLGQGSFRPSQQNPQDQGSVQPQQLPQFEEIRNLALQTLPAMCNVYIPPYC--TI
TraesCS6B02G065800-6C     QQQQLQQQQQQQQQQQQPSSQVSFQQPQQQYPSSQVSFQPSQLNPQAQGSVQPQQLPQFAEIRNLALQTLPAMCNVYIPPHCSTTI
TraesCS6B02G065856-6C     QQLQQQQQQQLHQQRQQPSSQVSFQQPQQQYPSSQVSFQPSQLNPQAQGSVQPQQLPQFAEIRNLALQTLPAMCNVYIPPHCSTTI
TraesCS6B02G086500-6C     QQQQQQQQQQQQQQQQQPSSQVSFQQPQQQYPSGQGSFQPSQQNPQAQSSVQHQQLPQFEEIRNLALQTLPAVCNVYIPPYCSTTI

                          :..350...
TraesCSU02G208300-7C      APVGIFGTN
TraesCSU02G220200-7C      APVGIFGTN
TraesCSU02G220600-7C      APVGIFGTN
TraesCSU02G108100-6C      APFGIFGTN
TraesCSU02G108400-6C      APVGFFGTN
TraesCSU02G108700-6C      APFGIFGTN
TraesCSU02G188800-6C      APVGIFGTN
TraesCSU02G108500-6C      AQVGIFGTN
TraesCSU02G239000-6C      APVGIFGTN
TraesCS6A02G048900-6C     APFGIFGTN
TraesCS6A02G049100-6C     APFGIFGTN
TraesCS6A02G049200-6C     VPFGIFGTN
TraesCS6A02G049400-6C     APFGIFGTN
TraesCS6A02G049600-6C     APFGIFGTN
TraesCS6A02G049700-6C     APFGIFGTN
TraesCS6A02G049800-6C     APFGIFGTN
TraesCS6B02G065800-6C     APFGIFGTN
TraesCS6B02G065856-6C     APFGIFGTN
TraesCS6B02G086500-6C     APFGIFGTN
```

**Fig. S6.** Diagnostic peptides for the 7-CYS α-gliadins detected in the proteomics study.

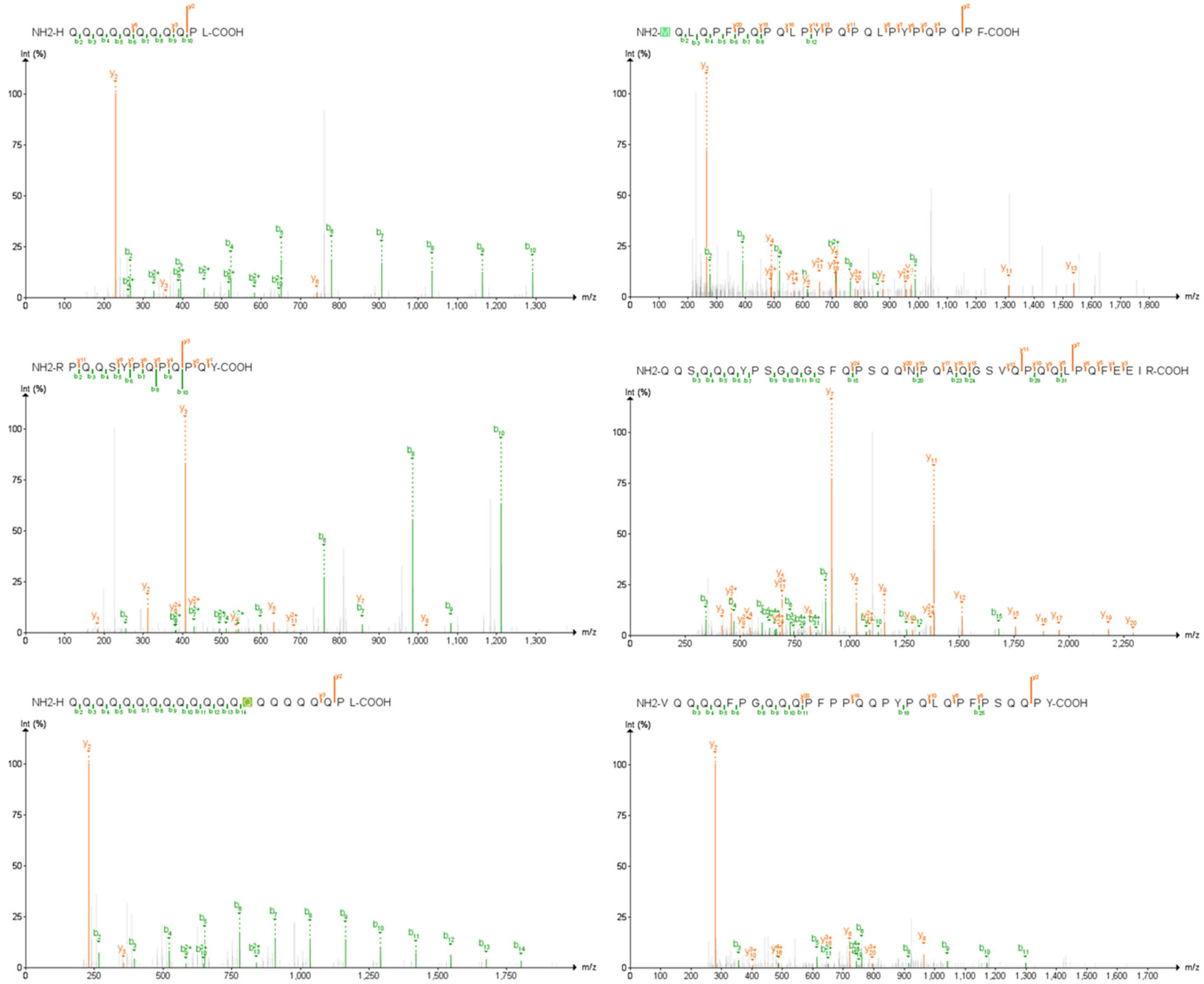
